# patternize: An R package for quantifying color pattern variation

**DOI:** 10.1101/121962

**Authors:** Steven M. Van Belleghem, Riccardo Papa, Humberto Ortiz-Zuazaga, Frederik Hendrickx, Chris D. Jiggins, W. Owen McMillan, Brian A. Counterman

**Affiliations:** Department of Zoology, University of Cambridge, Cambridge CB2 3EJ, United Kingdom; Department of Biology, Center for Applied Tropical Ecology and Conservation, University of Puerto Rico, Rio Piedras Campus, Puerto Rico; Department of Biological Sciences, Mississippi State University, 295 Lee Boulevard, Mississippi State, MS 39762, USA; Molecular Sciences and Research Center, University of Puerto Rico, San Juan, 00926, Puerto Rico.; Department of Computer Science, University of Puerto Rico, Rio Piedras Campus, Puerto Rico; Terrestrial Ecology Unit, Biology Department, Ghent University, Gent, Belgium; Royal Belgian Institute of Natural Sciences, Brussel, Belgium; Smithsonian Tropical Research Institute, Apartado 0843-03092, Panamá, Panama

**Keywords:** color patterns, heatmap, landmarks, image registration, image segmentation

## Abstract

1. The use of image data to quantify, study and compare variation in the colors and patterns of organisms requires the alignment of images to establish homology, followed by color-based segmentation of images. Here we describe an `R` package for image alignment and segmentation that has applications to quantify color patterns in a wide range of organisms.
2. `patternize` is an `R` package that quantifies variation in color patterns obtained from image data. `patternize` first defines homology between pattern positions across specimens either through manually placed homologous landmarks or automated image registration. Pattern identification is performed by categorizing the distribution of colors using an RGB threshold, *k*-means clustering or watershed transformation.
3. We demonstrate that `patternize` can be used for quantification of the color patterns in a variety of organisms by analyzing image data for butterflies, guppies, spiders and salamanders. Image data can be compared between sets of specimens, visualized as heatmaps and analyzed using principal component analysis (PCA).
4. `patternize` has potential applications for fine scale quantification of color pattern phenotypes in population comparisons, genetic association studies and investigating the basis of color pattern variation across a wide range of organisms.

## Introduction

Natural populations often harbor great phenotypic diversity. Variation in color and pattern are of the more vivid examples of morphological variability in nature. Taxa as diverse as spiders (De Busschere *et al.* 2012; Cotoras *et al.* 2016), insects (Katakura *et al.* 1994; Williams 2007), fish (Endler 1983; Houde 1987), amphibians and reptiles (Calsbeek *et al.* 2008; Allen *et al.* 2013; Balogová & Uhrin 2015; Rabbani *et al.* 2015), mammals (Hoekstra *et al.* 2006; Nekaris & Jaffe 2007) and plants (Clegg & Durbin 2000; Mascó *et al.* 2004) display natural variation in pigment or structural colorations. The distribution of colors in specific patterns play an important role in mate preference (Endler 1983; Kronforst *et al.* 2006), thermal regulation (Forsman *et al.* 2002), aposematism (Rojas *et al.* 2015) and crypsis (Nosil & Crespi 2006) and represent evolutionary adaptations that in many cases have promoted diversification within lineages.

Measuring phenotypic variation in organismal color patterns can provide insights into their underlying developmental and genetic architecture (Klingenberg 2010). However, precisely quantifying color pattern variation is challenging. Consistent comparisons of color patterns from images requires the (1) homologous alignment and (2) color-based segmentation of the images. Homologous alignment can be performed by transforming one image onto another. This transformation can be obtained from manually placed homologous landmarks or advanced image registration techniques, which can be stored and utilized to align color patterns extracted from the images. Image segmentation concerns the categorization of pixels by color. Previously, examples of color pattern quantification have been extensively developed for *Heliconius* butterflies (Color Pattern Modelling (CPM) in Le Poul *et al.* 2014) and primates (Allen *et al.* 2015). However, these applications are currently not easily accessible for use in other organisms. Similarly, advanced solutions are available for biomedical image analysis (Modat *et al.* 2010a; Schindelin *et al.* 2012, 2015), but are not tailored towards quantifying color pattern variation.

Here, we present `patternize`, an approach to quantification of color pattern variation from 2D images using the R statistical computing environment (R Development Core Team 2013). The package provides utilities to extract, transform and superimpose color patterns as well as downstream analysis (Fig. 1). The provided R functions combine single transformation and color extraction approaches. While transformations are obtained from manually placed homologous landmarks (`patLanRGB()`, `patLanK()` or `patLanW()`) or automated image registration `(patRegRGB()`, `patRegK()` or `patRegW()`), color-based segmentation of the patterns is performed by using threshold RGB (Red, Blue and Green) values (`patLanRGB()` or `patRegRGB()`), unsupervised classification of pixels into a set of clusters (`patLanK()` or `patRegK()`) or watershed transformation (`patLanW()` or `patRegW()`). By extracting and aligning color patterns from image data of large numbers of samples, p a t t e rn i z e provides quantitative measures of variation in color patterns that can be used for population comparisons, genetic association studies and investigating dominance and epigenetic interactions of color pattern variation in a wide range of organisms. We demonstrate the utility of the package with *Heliconius* butterflies and more challenging examples from guppy fish, Galápagos wolf spiders and salamanders.

**Fig. 1.**
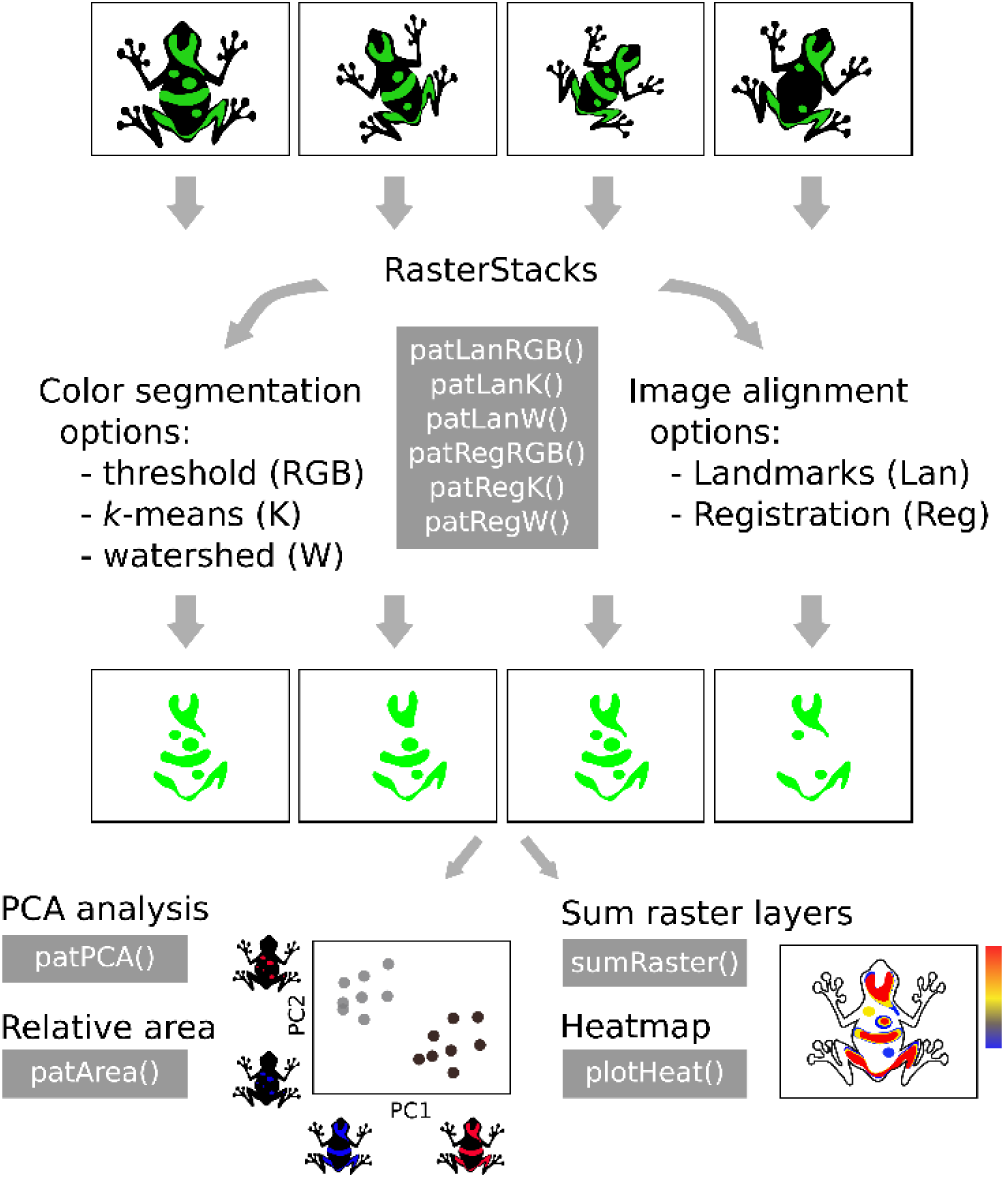
Overview of main `patternize` functions and functionality. Images can be aligned using homologous landmarks (Lan) or automatic registration (Reg), which aligns images using common intensity patterns. Colors can be extracted using an RGB threshold (RGB), *k*-means clustering (K) or by identifying watershed lines (W). The resulting extracted patterns can be summed and visualized as heatmaps or used for PCA analysis and calculating the relative area of the color patterns.

## Alignment

Superimposing color patterns to quantify variation in their expression requires the homologous alignment of the anatomical structures they occur in. Image transformations for this alignment can be obtained from landmark based transformations or image registration techniques.

### Landmark based transformations

Landmark based transformations use discrete anatomical points that are homologous among individuals in the analysis. Non-rigid, but uniform transformations from one set of ‘source’ landmarks to a set of ‘target’ landmarks such as *affine* transformations include translation, rotation, scaling and skewing (Hazewinkel 2001). Additionally, non-uniform changes in shape between the source and target landmarks can be accounted for by storing the transformation as if it were ‘the bending of a thin sheet of metal’, the so-called *thin plate spline* (TPS) transformation (Duchon 1976). Both the affine and TPS transformation can be calculated from sets of landmarks (Fig. 2A). We implemented these landmark transformations using utilities provided by the R package `Morpho` (Schlager 2016). Landmarks can be transformed using an arbitrarily chosen reference sample or an average landmark shape obtained from a set of samples. The average landmark shape is obtained by means of Procrustes superimposition of the samples (Goodall 1991).

**Fig. 2.**
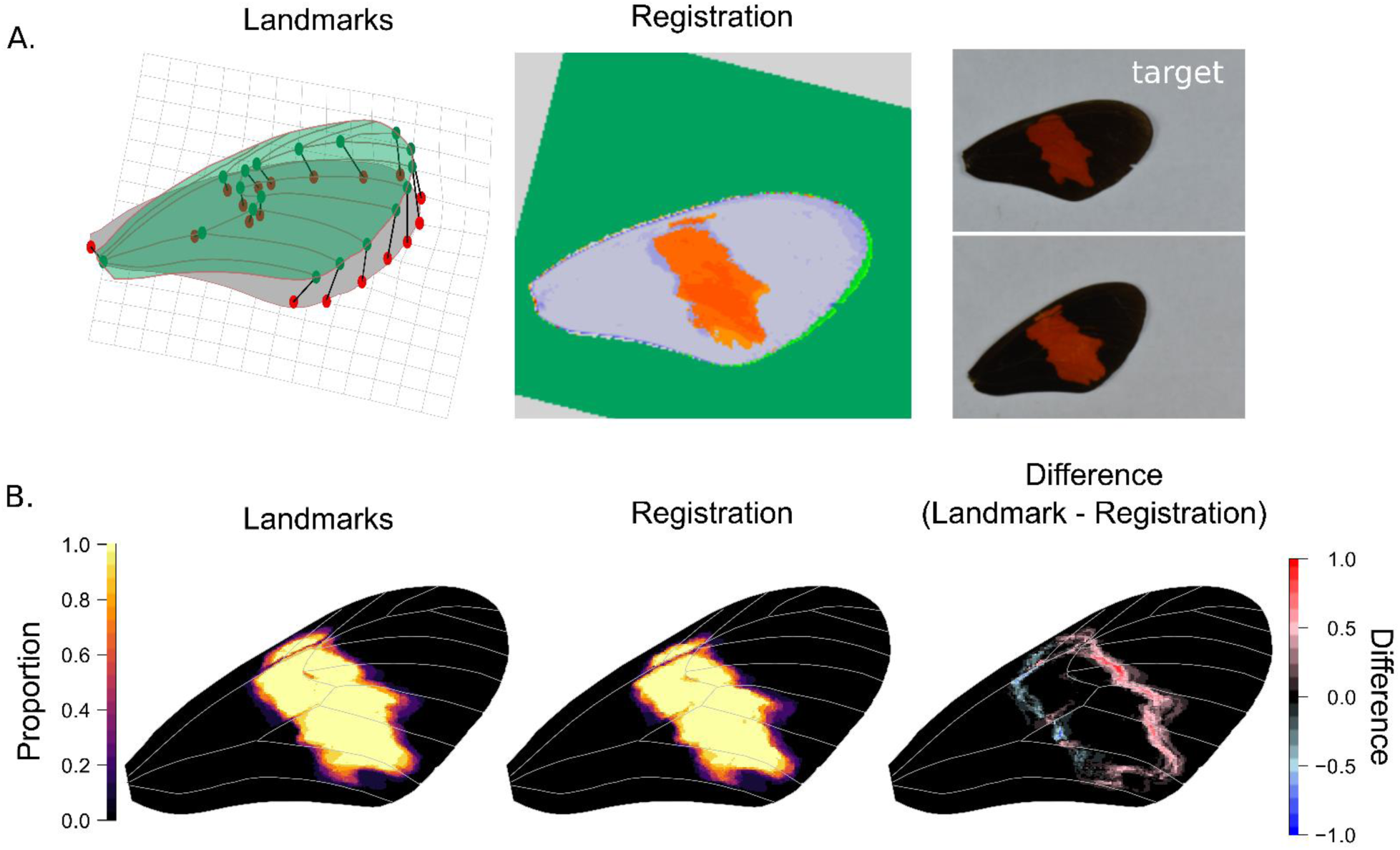
Comparison of image transformation using landmarks or automated registration for quantification of color pattern variation. (A.) Illustration of transformation strategies of a source (green) image to a target (gray) image. The thin plate spline (TPS) transformation from the source to target landmarks is illustrated by the deformed grid and can be used to transform the image or extracted color pattern. Image registration attempts to find common patterns in images and align the source (green) image to the pixel coordinate system of the target (gray) image. Note the extracted color pattern in red. (B.) Example comparison between landmark approach for color pattern alignment for ten butterfly wings of male *Heliconius erato hydara*. For the landmark approach, we used TPS transformation. For the image registration approach, we used affine transformation and 75% of sub-volumes included as inliers.

### Image registration

Alternative to landmark based methods, fast and accurate image registration techniques are available for calculating a transformation from a source to target image based on either intensity patterns or features such as points, lines or contours present in the images (Goshtasby 2005) (Fig. 2A). We use a computation efficient intensity-based image registration technique implemented in the NiftyReg image registration library (Translational Imaging Group (TIG) 2016) and made available in R through the `RNiftyReg` package (Clayden *et al.* 2017). This methodology calculates the global transformation of an image by finding correspondences between sub-volumes of the two images (Modat *et al.* 2010a,b). Correspondence is assessed using intensity-based similarity measures and used to calculate the transformation parameters through a least trimmed square (LTS) regression method (Modat *et al.* 2010a,b). The number of corresponding sub-volumes to be included or considered as outliers in the calculation of the transformation can be varied by the user. The global transform calculated by NiftyReg can be rigid (i.e. including translation, rotation and scaling) or affine (i.e. translation, rotation, scaling and skewing).

## Color pattern extraction

Studying variation in color patterns requires the correct identification of the color boundaries. `patternize` provides functionality to categorize the distribution of colors using either an RGB threshold, *k*-means clustering or watershed transformation.

### RGB threshold

Color boundaries can be extracted from images or the trait of interest using an RGB threshold (Fig. 2 and Fig. 3). By selecting pixels within a specified color range (specified as RGB value and offset) we provide a basic image segmentation approach that works well for extracting distinct color patterns. Additionally, for distinct color patterns, the RGB value can be iteratively recalculated as the average for the extracted color pixels. This latter approach permits patterns to be easily combined when extracted from sets of images that may have been taken under different light conditions resulting in differences in intensity and contrast.

**Fig. 3.**
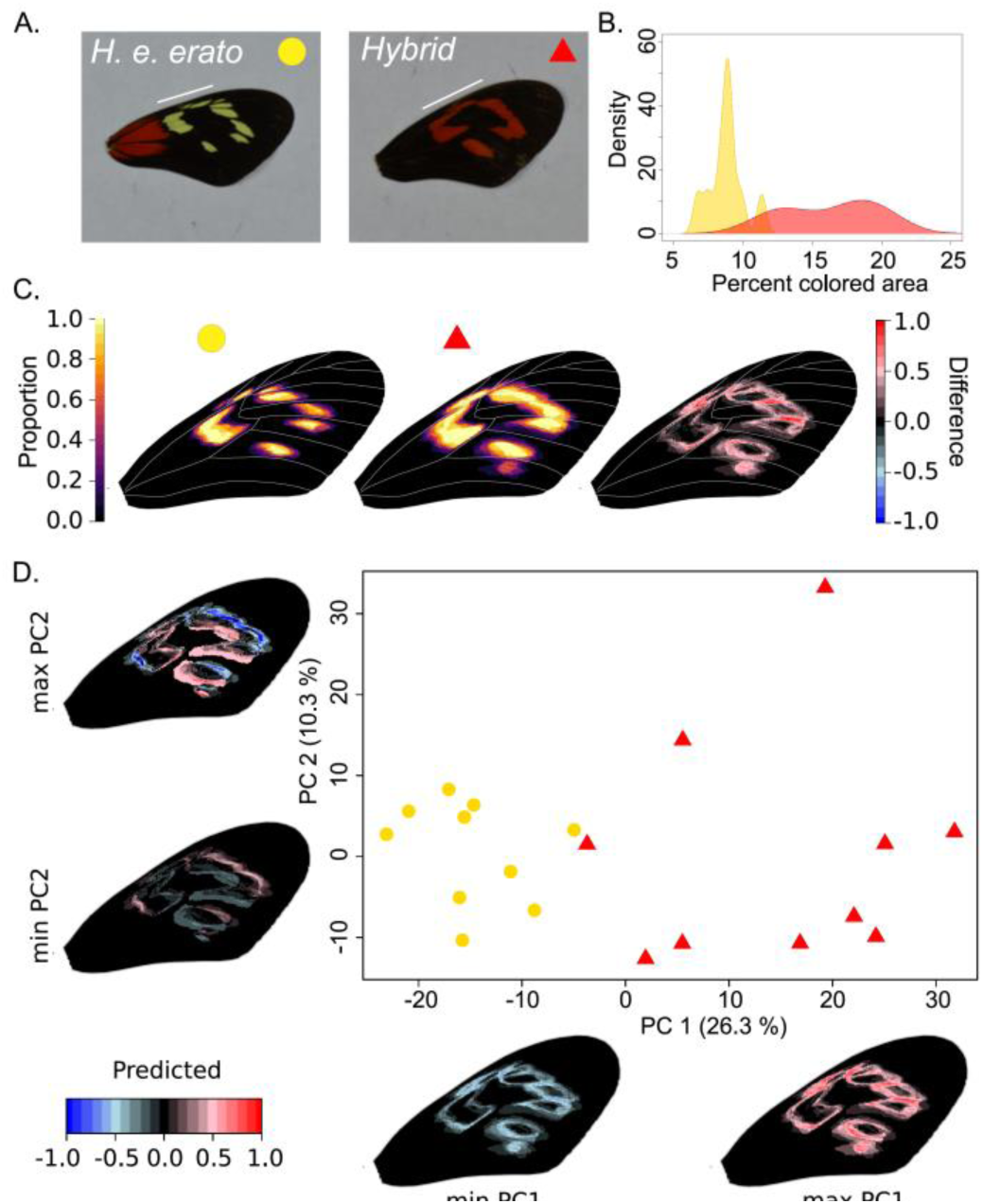
Example of image registration and threshold color extraction in the forewing band area of *Heliconius erato erato* (n = 10) and hybrid (n = 10) butterflies (French Guiana). (A.) Example of original images with a white line indicating the forewing band area. The hybrid represents a naturally occurring backcross in a hybrid zone with *H. e. hydara* (see Fig. 1) that results in red color expression in the forewing band. (B.) Density plot showing the probability to find a sample with a certain percentage of colored area in the wing expressing yellow in *H. e. erato* and red in the hybrid. (C.) Visualizing the variation in color pattern expression in a heatmap shows a consistently larger pattern in the hybrid phenotypes (*H. e. erato*: left, hybrid: middle, hybrid minus *H. e. erato*: right). (D.) Principal component analysis (PCA) confirms that the main axis of variation (PC1) is related to size of the pattern (yellow or red in *H. e. erato* and hybrids, respectively) and separates the *H. e. erato* and hybrid samples. The second principal component (PC2) axis highlights more complex shape differences in the forewing band among the samples as demonstrated by the shape changes of the color patterns along the principal component axis.

### *k*-means clustering

We implemented an unsupervised approach for color-based image segmentation by using *k*-means clustering (Fig. 4 and Fig. 5) (Hartigan & Wong 1979). This algorithm assigns pixel RGB values to *k* clusters by iteratively assigning each pixel in the image to the RGB cluster that minimizes the distance between the pixel and the cluster centers. Cluster centers are recalculated each iteration by averaging all pixels in the cluster until convergence. We implemented *k*-means clustering using the R package `stats` (R Development Core Team 2013).

**Fig. 4.**
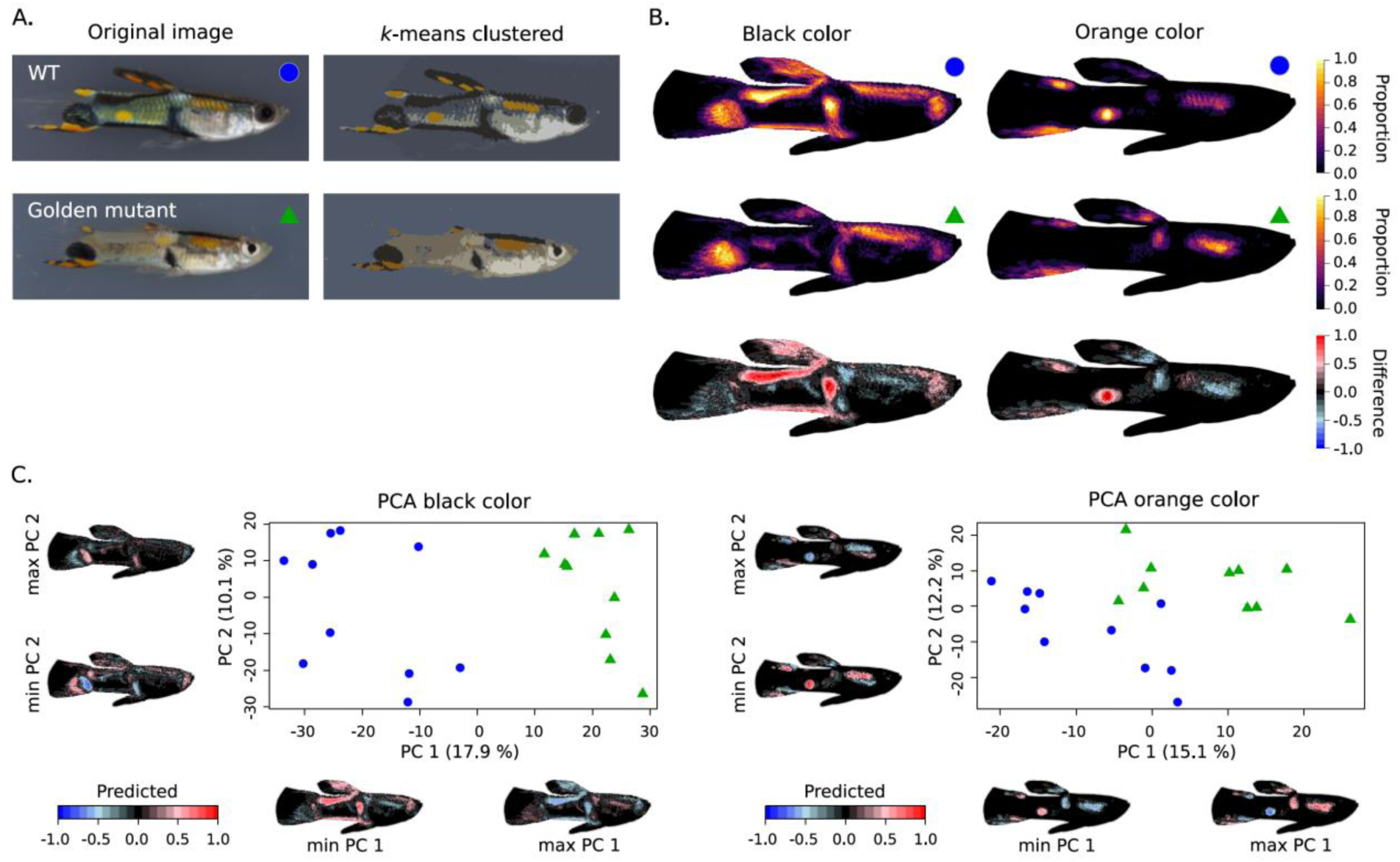
Example of image registration and *k*-means clustering of colors in guppies (*Poecilia reticulata*). (A.) Original image of a wild type (WT) and *golden* mutant guppy and their *k*-means clustered representation (clusters = 7). (B.) Heatmaps and difference between WT (n=10) and golden mutant (n=10) for black and orange color clusters. (D.) PCA analysis of the pixel matrices obtained for the black (left) and orange (right) color clusters. Images were obtained with permission from Kottler *et al.* (2013).

**Fig. 5.**
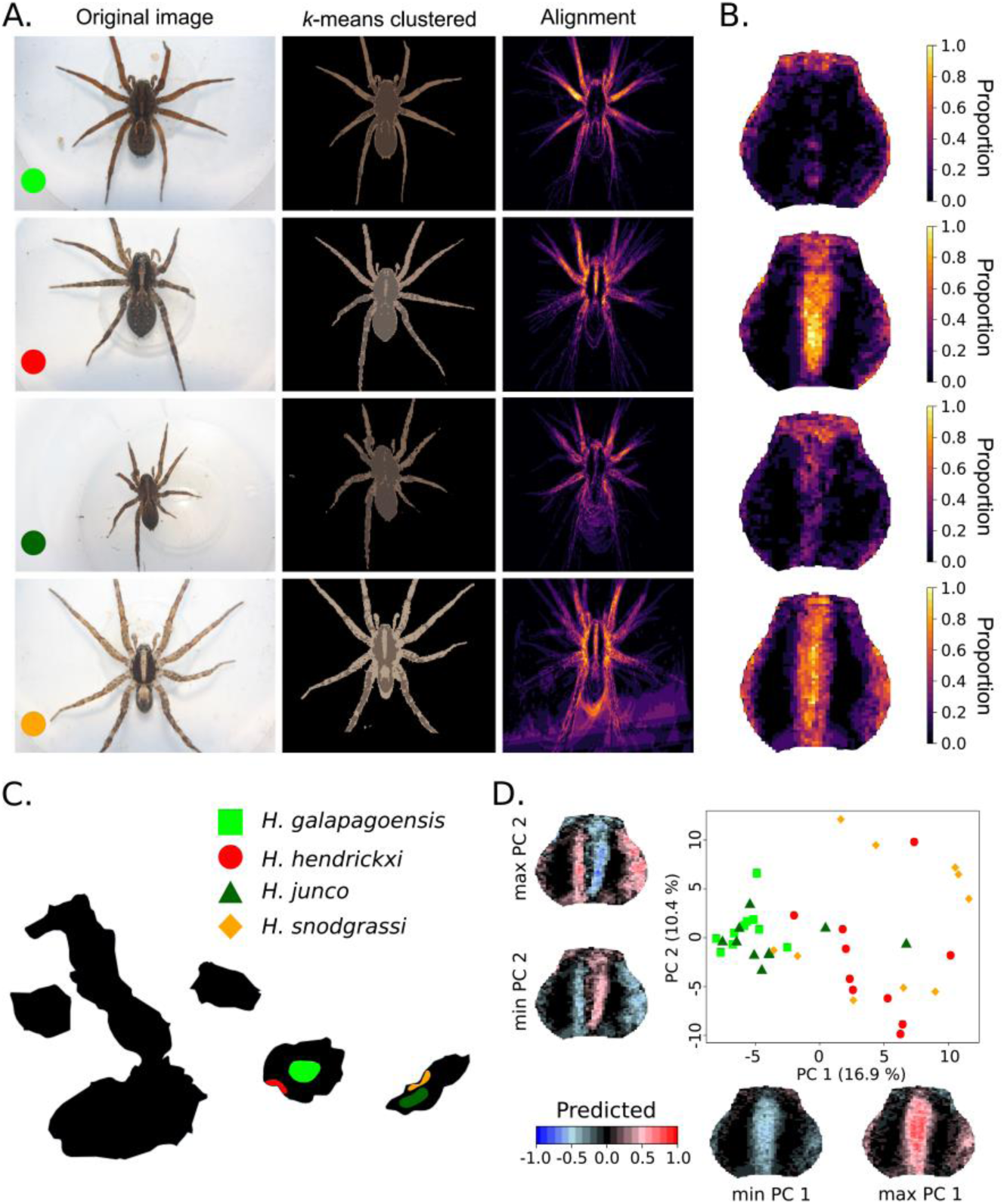
Example of image registration and *k*-means clustering of the color pattern of Galápagos wolf spiders (*Hogna*). (A.) From left to right: example of original image (10 images were used for each species), *k*-means clustered image (k = 3) with removed background, and alignment of the lightest color. (B.) Heatmap corresponding to the lightest color cluster focused on the carapace. (C.) Map of the Galápagos islands with colors indicating the distribution of four *Hogna* species, two high elevation species (light and dark green) and two coastal species (red and orange). (D.) PCA analysis of the pixel matrices obtained for the lightest color cluster demonstrates that the coastal (*H. hendrickxi* and *H. snodgrassi*) and high-elevation (*H. galapagoensis* and *H. junco*) species cluster phenotypically together and share, respectively, the presence and absence of a pale median band on their carapace. Images were obtained with permission from De Busschere *et al.* (2012).

Clusters are first obtained from a reference image and then used as initial cluster centers for the *k*-means clustering of the subsequently analyzed images. This allows the program to match clusters that represent the same color pattern in different images. For *k*-means clustering, the number of clusters must be defined manually. For organisms with less distinct pattern boundaries, this is best done by testing different numbers of clusters and choosing a number that best assigns pixels to color patterns.

### Watershed transformation

The watershed transformation is a powerful tool for image segmentation (Fig. 6) (Beucher 1991). The concept of watershed treats the image as a topographic map by calculating a gradient map with high values in parts of the image where pixel values change abruptly (Fig. 6B). Subsequently, a flooding process propagates pattern and background labels guided by the gradient map. Continuing the flooding until pattern and background labels meet, determines the watershed lines (ridges in the topography) that are used to segment the image (Fig. 6C). We implemented the watershed algorithm with utilities from the R package `imager` (Barthelme 2017) that is based on the image processing library CImg (Tschumperle 2004). In our implementation, the pattern and background labels are chosen by manually identifying at least one pattern and one background pixel (at least one for each separate pattern and background element). This manual assignment helps the user to overcome potential differences in image lightning, glare or overlap between pattern and background RGB values.

**Fig. 6.**
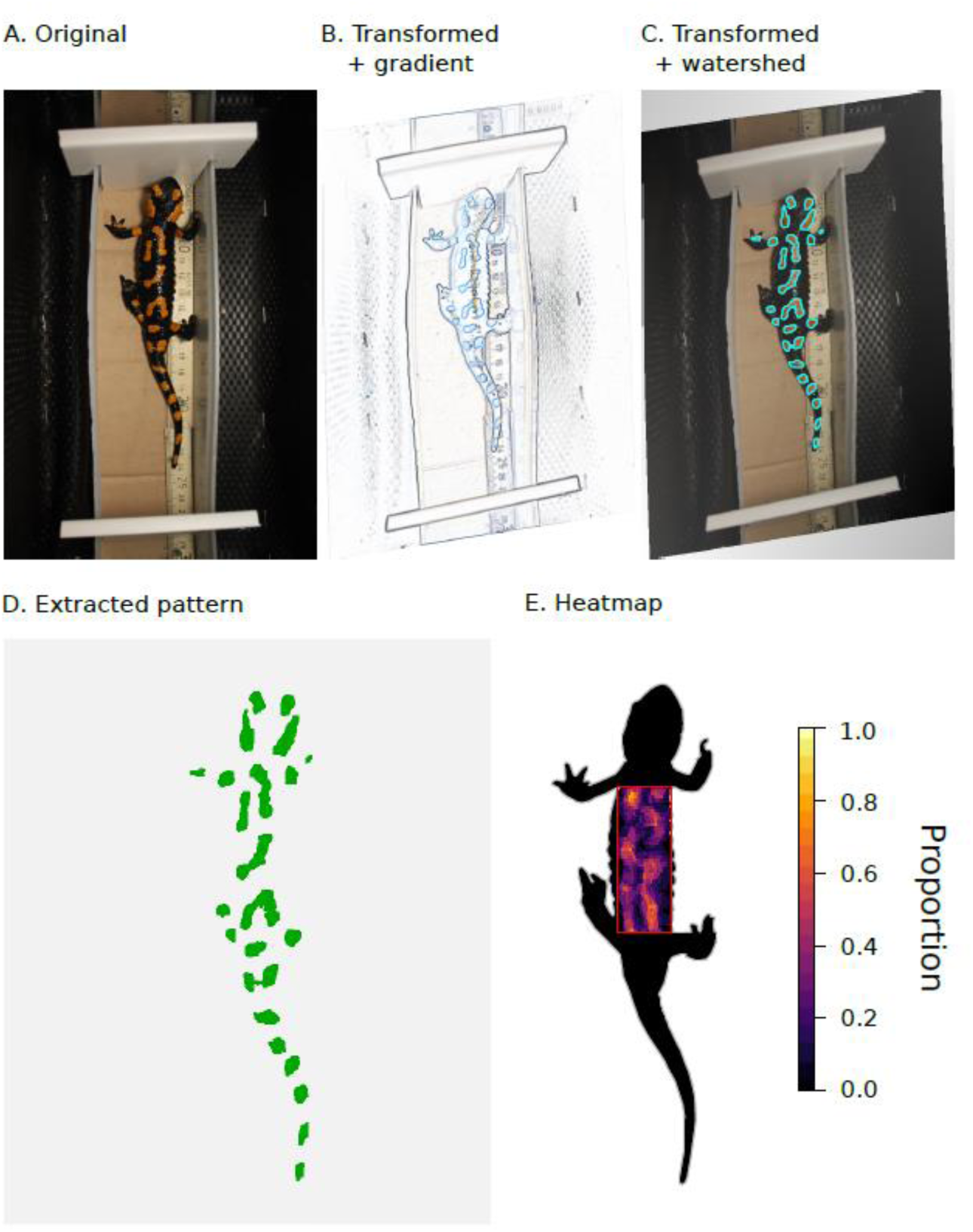
Example of watershed transformation for color pattern extraction in fire salamanders (*Salamandra salamandra*). (A.) Original image. (B.) Image gradient transformed to a reference shape using landmarks. (C.) Transformed image with watershed lines highlighted. (D.) Extracted patterns using the watershed lines. (E.) Heatmap of orange patterns extracted from ten male fire salamanders. Areas outside the red box were masked. Images were obtained with permission from Balogová & Uhrin (2015).

## Output

The main `patternize` functions generate a list of extracted color patterns from each image stored as a `raster` object (Hijmans 2016). These extracted patterns can be summed and visualized as heatmaps or used to calculate the relative area of the color patterns. To better characterize variation in color patterns among samples, we implemented linear principal component analysis (PCA). For an extracted color pattern, PCA can be performed on the binary representation of the aligned color pattern rasters obtained from each sample (Fig. 3-5). In this matrix, pixel coordinates that have the color of interest in a sample have a value of one, whereas pixel coordinates without the color have the value zero assigned. The variance-covariance matrix obtained from the binary matrix for a color is suitable for PCA, which allows visualizing the main variations in color pattern boundaries among or between groups of samples, as well as the predicted color pattern changes along the principal component (PC) axis (Johnson & Wichern 2007). In the visualization of the predicted color pattern changes, positive values present a higher predicted expression of the pattern, whereas negative values present the absence of the pattern. Note that parts of the color patterns that are expressed in all considered samples have a predicted value of zero, as these pixels do not contribute variance for the PCA analysis.

## Examples

### RGB threshold pattern extraction in *Heliconius* butterflies

We demonstrate the utility of image alignment and RGB threshold extraction in the forewing band area of *Heliconius erato* populations (Fig. 2). *Heliconius* butterflies from the Neotropics display great diversity in forewing band shape, which is mainly defined by expression of the *wntA* gene (Van Belleghem *et al.*; Martin *et al.* 2012). Expression of red pigments in the wing scales is on its turn defined by expression of the *optix* gene (Reed *et al.* 2011). Comparison of the landmark and image registration approach applied to the red forewing band variation in *H. erato hydara* shows that both methods perform well (Fig. 2B). The TPS transformation used in the landmark approach resulted in a better fit to the internal structures of the wing (i.e. wing veins). The slight offset between the color pattern and vein position in the image registration approach likely resulted from a bias in the linear transformation towards aligning the outline of the wing and not including non-uniform changes in shape within the wing.

Next, we performed automated image registration and RGB threshold color extraction on the same forewing band area of *H. erato erato*. In this region of the wing, *H. e. erato* lacks *optix* expression and, thus, red scales. However, naturally occurring hybrids between *H. e. erato* and *H. e. hydara* show *optix* expression in the forewing band area (Fig. 3). With this example, we demonstrate the ability to compare homologous, but differing colored pattern elements (i.e. yellow versus red). The PCA analysis and relative area of the extracted patterns allow to differentiate the two groups of butterflies and indicate overexpression of the color pattern in hybrids.

### Automated registration and k-means clustering in guppies and spiders

To assess the general utility of our application across taxa, we applied the automated registration and *k*-means clustering approach to groups with more complex body shape and color pattern variation; guppy fish and Galápagos wolf spiders. Males of the guppy (*Poecilia reticulata*) vary greatly in their ornamental patterns that have evolved in response to both natural and sexual selection. Several mutants have been described among male guppies that affect color pattern expression. Manually quantifying the differences in color pattern expression among these mutations has provided valuable insights into the developmental basis and interactions of the involved genes (Kottler *et al.* 2013). Here, we summarized and compared the black and orange color patterns expressed in wild type (WT) versus *golden* mutants of *P. reticulata* males using images obtained from Kottler *et al.* 2013 (images were used from backcrosses obtained from *golden blue* mutant females with heterozygous males from crossing *golden blue* with inbred wild-derived Cumána populations) (Fig. 4). All images were aligned to a target image using image registration and colors were *k*-means clustered into seven groups. Before *k*-means color clustering, the background was masked using the outline of the guppy in the target image. Our analysis of the black and orange color cluster strongly matched the description presented in Kottler *et al.* 2013, demonstrating the absence of a posterior orange spot in *golden* mutants backcrossed into a Cumána population genetic background and more diffuse and shifted black ornaments in the *golden* mutants.

Wolf spiders of the genus *Hogna* inhabit high elevation and coastal habitats on the Galápagos islands Santa Cruz and San Cristobal (De Busschere *et al.* 2010). Despite the phylogenetically close relationship of the high elevation and coastal populations within both islands, morphometric analysis, including measurements of color intensity, have highlighted striking parallel phenotypic divergence between the high elevation and coastal species between the islands (De Busschere *et al.* 2012). Coastal species appear to be paler with a more conspicuous median band on the carapace compared to high elevation species. Here, we demonstrate the robustness of automated image registration by aligning the highly variable images of the wolf spiders (Fig. 5). By focusing on correspondence between the images, the automated image registration technique manages to align the spider’s carapace, which is morphologically the most consistent part in the images. By assigning colors in the spiders to only two clusters, we show a similar pattern as described in De Busschere *et al.* (2012) in which the coastal species show a consistently broader and more conspicuous median band on the carapace and pale lateral bands compared to the high elevation species.

### Watershed pattern extraction in fire salamanders

The glare that is usually present in images of amphibians can make it challenging to correctly extract the color patterns. Additionally, some pattern elements may be difficult to identify based on color alone. To overcome these difficulties, we illustrate the watershed segmentation using images of fire salamanders obtained from Balogová & Uhrin (2015) (Fig. 6). The fire salamander (*Salamandra salamandra*) is common to Europe and is black with yellow, orange or red spots or stripes. The watershed approach confidently identifies the orange pattern boundaries in the analyzed images. Combining this color pattern extraction approach with aligning the images allows users to identify regions in the salamander’s body where spots or stripes are more consistently expressed.

## Concluding remarks and recommendations

### Alignment

`patternize` provides an unbiased, fast and user-friendly approach for color pattern analysis that is applicable to a wide variety of organisms. `patternize` takes jpeg images as input, which can be downsampled to decrease computation times. While the landmark based approach is computationally slightly faster, automated image registration removes the need for labor-intensive landmark setting. Moreover, image registration reduces any variation introduced by differences in how users manually place image landmarks. However, because automated registration uses intensity patterns in the images, it can be highly sensitive to artifacts in the background and care should be taken by standardizing the experimental setup. For cases in which the background differs starkly from the studied object, functionality is included that allows users to remove the background by providing RGB cutoff values. The package also allows users to review the image registration progress to assess the quality of the automatic registration.

### Color pattern extraction

Variation in photographic conditions complicates color pattern extraction. The option for iteratively recalculating the RGB value and defining the start clusters for k-means clustering from a reference image can improve color pattern extraction under these conditions. However, setting correct RGB or cluster parameters may impact results and should be optimized for each analysis. Appropriate RGB and offset values can be obtained, for instance, by extracting RGB values from image pixels or areas of interest (e.g. use `sampleRGB ()`). Using few or many *k*-means clusters may, respectively, result in grouping colors of interest or assigning multiple clusters to a single pattern of interest. Finally, in contrast to RGB threshold color extraction and *k*-means clustering, watershed transformation takes into account the spatial proximity of pixels. Doing so, the interactive identification of pattern versus background in the watershed transformation provides a way to extract color patterns that is robust to variation in photographic conditions.

### Output

The output of the main `patternize` functions are `raster` objects (Hijmans 2016) that provide for a wide range of downstream analyses. As demonstrated by the examples, we provide functions to intersect (mask) the extracted patterns with defined outlines, sum or subtract the patterns to plot heatmaps, calculate the relative area in which the pattern is expressed and carry out principal component analysis (PCA). Overall, we hope this R package provides a useful tool for the community of researchers working on color and pattern variation in animals.

## Acknowledgments

We kindly thank Jon Clayden for help with implementing RNiftyReg, Stefan Schlager for help with Morpho, Maria Bencomo, Emily Shelby and Heather Smith for help with digitizing the *Heliconius* images and Verena Kottler, Charlotte De Busschere and Monika Balogová for allowing us to use the guppy, wolf spider and fire salamander images, respectively. SMVB and BAC were funded by NSF grant DEB-1257839. HOZ was supported in part by NIH grant 5P20GM103475- 13. All authors declared that they have no conflict of interest.

## Data accessibility

The package and descriptions of the functions and parameters are available as library(“patternize”) on CRAN (cran.r-project.org/web/packages/patternize). The code, ongoing developments and data and code used for the examples can be accessed through GitHub (github.com/StevenVB12/patternize; github.com/StevenVB12/patternize-examples). Bug reports and feature requests can be sent using the GitHub issue tracker.

## Author contributions

SMVB and BAC conceived the development of the package. SMVB wrote the code. SMVB, RP and BAC wrote the manuscript. HOZ helped improving the code. SMVB, BAC, FH and RP conceived data acquisition. HOZ, FH, CDJ and WOM contributed helpful comments for building the package and writing the manuscript. All authors contributed critically to the drafts and gave final approval for publication.

## References

Allen, W.L., Baddeley, R., Scott-samuel, N.E. & Cuthill, I.C. (2013). The evolution and function of pattern diversity in snakes. Behavioral Ecology, 24, 1237–1250.

Allen, W.L., Higham, J.P. & Allen, W.L. (2015). Assessing the potential information content of multicomponent visual signals: a machine learning approach. Proceedings of the Royal Society B, 282, 20142284.

Balogová, M. & Uhrin, M. (2015). Sex-biased dorsal spotted patterns in the fire salamander (*Salamandra salamandra*). Salamandra, 51, 12–18.

Barthelme, S. (2017). imager: Image processing library based on ‘CImg’. R-package version 0.40.2. https://CRAN.Rproject.org/package=imager.

Beucher, S. (1991). The watershed transformation applied to image segmentation. Proc. 10th Pfefferkorn Conf. on Signal and Image Processing in Microscopy and Microanalysis, pp. 299–314.

Calsbeek, R., Bonneaud, C. & Smith, T.B. (2008). Differential fitness effects of immunocompetence and neighbourhood density in alternative female lizard morphs. 103–109.

Clayden, J., Modat, M., Presles, B., Anthopoulos, T. & Daga, P. (2017). RNiftyReg: Image Registration Using the ‘NiftyReg’ Library. R package version 2.5.0. https://CRAN.R-project.org/package=RNiftyReg.

Clegg, M.T. & Durbin, M.L. (2000). Flower color variation: A model for the experimental study of evolution. Proceedings of the National Academy of Sciences, 97, 7016–7023.

Cotoras, D.D., Brewer, M.S., Croucher, P.J.P., Geoff, S., Lindberg, D.R. & Gillespie, R.G. (2016). Convergent evolution in the colour polymorphism of *Selkirkiella* spiders (Theridiidae) from the South American temperate rainforest. Biological Journal of the Linnean Society.

De Busschere, C., Baert, L., Van Belleghem, S.M., Dekoninck, W. & Hendrickx, F. (2012). Parallel phenotypic evolution in a wolf spider radiation on Galápagos. Biological Journal of the Linnean Society, 106, 123–136.

De Busschere, C., Hendrickx, F., Van Belleghem, S.M., Backeljau, T., Lens, L. & Baert, L. (2010). Parallel habitat specialization within the wolf spider genus *Hogna* from the Galápagos. Molecular ecology 19, 4029–4045.

Duchon, J. (1976). Splines minimizing rotation invariant semi-norms in Sobolev spaces. Volume 571 of the series Lecture Notes in Mathematicss (eds W. Schempp & K. Zeller), pp. 85–100. Springer.

Endler, J.A. (1983). Natural and sexual selection on color patterns in poeciliid fishes. Environmental Biology of Fishes, 9, 173–190.

Forsman, A., Ringblom, K., Civantos, E. & Ahnesjö, J. (2002). Coevolution of color pattern and thermoregulatory behavior in polymorphic pygmy grasshoppers *Tetrix undulata*. Evolution, 56, 349–360.

Goodall, C. (1991). Procrustes methods in the statistical analysis of shape. Journal of the Royal Statistical Society. Series B, 53, 285–339.

Goshtasby, A. (2005). 2-D and 3-D image registration: for medical, remote sensing, and industrial applications. Wiley, Hoboken, NJ.

Hartigan, J.A. & Wong, M.A. (1979). Algorithm AS 136: A k-means clustering algorithm. Journal of the Royal Statistical Society. Series B, 28, 100–108.

Hazewinkel, M. (Ed.). (2001). Affine transformation. Encyclopedia of Mathematics. Springer.

Hijmans, R.J. (2016). raster: Geographic data analysis and modeling. R package version 2.5-8. http://cran.rproject.org/package=raster.

Hoekstra, H.E., Hirschmann, R.J., Bundey, R. a, Insel, P. a & Crossland, J.P. (2006). A single amino acid mutation contributes to adaptive beach mouse color pattern. Science, 313, 101–104.

Houde, A.E. (1987). Mate choice based upon naturally occurring color-pattern variation in a guppy population. Evolution, 41, 1–10.

Johnson, R.A. & Wichern, D.W. (2007). Applied multivariate statistical analysis, 6th Edition. Pearson.

Katakura, H., Saitoh, S., Nakamura, K. & Abbas, I. (1994). Multivariate analyses of elytral spot patterns in the phytophagous ladybird beetle *Epilachna vigintioctopunctata* (Coleoptera, Coccinellidae) in the province of Sumatra Barat, Indonesia. Zoological science, 11, 889–894.

Klingenberg, C.P. (2010). Evolution and development of shape: integrating quantitative approaches. Nature, 11, 623–635.

Kottler, V.A., Fadeev, A., Weigel, D. & Dreyer, C. (2013). Pigment pattern formation in the guppy, *Poecilia reticulata*, involves the Kita and Csf1ra receptor tyrosine kinases. Genetics, 194, 631–646.

Kronforst, M.R., Young, L.G., Kapan, D.D., McNeely, C., O’Neill, R.J. & Gilbert, L.E. (2006). Linkage of butterfly mate preference and wing color preference cue at the genomic location of *wingless*. Proceedings of the National Academy of Sciences of the United States of America, 103, 6575–6580.

Le Poul, Y., Whibley, A., Chouteau, M., Prunier, F., Llaurens, V. & Joron, M. (2014). Evolution of dominance mechanisms at a butterfly mimicry supergene. Nature Communications, 5, 1–8.

Martin, A., Papa, R., Nadeau, N.J., Hill, R.I., Counterman, B.A., Halder, G., Jiggins, C.D., Kronforst, M.R., Long, A.D., McMillan, W.O. & Reed, R.D. (2012). Diversification of complex butter flywing patterns by repeated regulatory evolution of a Wnt ligand. Proceedings of the National Academy of Sciences of the United States of America, 109, 12632–12637.

Mascó, M., Noy-Meir, I. & Sérsic, A.N. (2004). Geographic variation in flower color patterns within *Calceolaria uniflora* Lam. in Southern Patagonia. Plant Systematics and Evolution, 244, 77–91.

Modat, M., Mcclelland, J. & Ourselin, S. (2010a). Lung registration using the NiftyReg package. Medical Image Analysis for the Clinic: A Grand Challenge, Workshop Proc. from MICCAI 2010, 33–42.

Modat, M., Ridgway, G.R., Taylor, Z.A., Lehmann, M., Barnes, J., Hawkes, D.J., Fox, N.C. & Ourselin, S. (2010b). Fast free-form deformation using graphics processing units. Computer Methods and Programs in Biomedicine, 98, 278–284.

Nekaris, K.A.I. & Jaffe, S. (2007). Unexpected diversity of slow lorises (*Nycticebus spp.*) within the Javan pet trade: implica- tions for slow loris taxonomy. Contributions to Zoology, 76, 187–196.

Nosil, P. & Crespi, B.J. (2006). Experimental evidence that predation promotes divergence in adaptive radiation. Proceedings of the National Academy of Sciences, 103, 9090–9095.

Rabbani, M., Zacharczenko, B., Green, D.M., Abbani, M.O.R. & Acharczenko, B.R.Z. (2015). Color pattern variation in a cryptic amphibian, *Anaxyrus fowleri*. Journal of Herpetology, 49, 649–654.

R Development Core Team. (2013). R: A language and environment for statistical computing. R Foundation for Statistical Computing, Vienna, Austria.

Reed, R.D., Papa, R., Martin, A., Hinas, H.M., Counterman, B.A., Pard-Diaz, C., Jiggins, C.D., Chamberlain, N.L., Kronforst, M.R., Chen, R., Nijhout, H.F. & McMillan, W.O. (2011). *optix* drives the repeated convergent evolution of butterfly wing pattern mimicry. Science, 333, 1137–1141.

Rojas, B., Valkonen, J. & Nokelainen, O. (2015). Aposematism. Quick guide. Current biology, 25, R350–R351.

Schindelin, J., Arganda-carreras, I., Frise, E., Kaynig, V., Longair, M., Pietzsch, T., Preibisch, S., Rueden, C., Saalfeld, S., Schmid, B., Tinevez, J., White, D.J., Hartenstein, V., Eliceiri, K., Tomancak, P. & Cardona, A. (2012). Fiji: an open-source platform for biological-image analysis. Nature Methods, 9, 676–682.

Schindelin, J., Rueden, C.T., Hiner, M.C. & Eliceiri, K.W. (2015). The ImageJ ecosystem: An open platform for biomedical image analysis. Molecular reproduction & Development, 82, 518–529.

Schlager, S. (2016). Morpho: Calculations and visualisations related to geometric morphometrics. R package version 2.3.1.1. http://cran.r-project.org/package=Morpho.

Translational Imaging Group (TIG). (2016). NiftyReg. https://sourceforge.net/projects/niftyreg/.

Tschumperle, D. (2004). The CImg library: http://cimg.sourceforge.net. The C++ Template Image Processing Library.

Van Belleghem, S.M., Rastas, P., Papanicolaou, A., Martin, S.H., Arias, C.F., Supple, M.A., Hanly, J.J., Mallet, J., Lewis, J.J., Hines, H.M., Ruiz, M., Salazar, C., Linares, M., Moreira, G.R.P., Jiggins, C.D., Counterman, B.A., McMillan, W.O. & Papa, R. Complex modular architecture around a simple toolkit of wing pattern genes. Nature Ecology & Evolution.

Williams, P. (2007). The distribution of bumblebee colour patterns worldwide: possible significance for thermoregulation, crypsis, and warning mimicry. Biological Journal of the Linnean Society, 92, 97–118.

